# A neuronal atlas of neurexin isoform expression in *Caenorhabditis elegans*

**DOI:** 10.64898/2026.11.14.766646

**Authors:** Elisa B. Frankel, Parise S. Henry, Nikos Karmirantzos, Maggi Leider, Peri T. Kurshan

## Abstract

The *Caenorhabditis elegans* neurexin gene *nrx-1* encodes both long (α) and short (γ) isoforms, but how their expression varies across individual neurons remains unclear. Using NeuroPAL and endogenous transcriptional reporters, we generated a nervous system-wide atlas of *nrx-1* expression in L4 hermaphrodites. We assessed reporters for one α and two γ transcript groups in 267 identified neurons. The two γ reporters collectively marked a broader set of neurons than the α reporter, and most neurons expressing the α reporter also expressed at least one γ reporter. Among the ventral nerve cord motor neurons examined, α reporter expression was restricted to cholinergic neurons, while one γ reporter was detected exclusively in GABAergic neurons. The other γ reporter was consistently detected in most cholinergic motor neurons and detected in only a subset of GABAergic motor neurons, often variably across animals. In two-fold embryos, this same γ reporter was detected in more neurons than the α reporter. Two upstream regulatory elements contributed to its expression in different neuronal populations. At dorsal nerve cord synapses, disruption of either isoform class reduced NRX-1 fluorescence, with a larger reduction after disruption of the short isoforms. By identifying neurons that express different combinations of long and short neurexins, this atlas provides a starting point for testing where the two isoform classes have distinct synaptic roles and where they may act together.

## INTRODUCTION

Neurexins are central to synapse development and function, but the mechanisms that generate their diverse effects across neuronal cell types remain unclear. Neurexins can regulate synapse organization, neurotransmitter release, postsynaptic receptor localization, and synaptic maintenance, but they do not perform each of these functions at all synapses (Südhof, 2017). In *C. elegans,* for example, presynaptic NRX-1 is required for the formation of cholinergic synapses onto GABAergic neurons but not neighboring muscle cells, even though both synaptic partners receive inputs from the same cholinergic motor neurons (Philbrook et al., 2018). Neurexin requirements also differ across neuronal populations in vertebrate nervous systems, suggesting that their functions depend on the cellular and synaptic context in which they operate (Ducrot et al., 2023; Jiang et al., 2026). These observations suggest that the context-dependent functions of neurexin arise, in part, from differences in isoform expression.

In mammals, three neurexin genes (*Nrxn1*, *2* and *3*) use multiple promoters and extensive alternative splicing to generate structurally different α and β isoforms (Südhof, 2017; Treutlein et al., 2014; Ullrich et al., 1995). The *Nrxn1* locus contains an additional internal promoter that produces the much shorter Nrxn1γ isoform (Sterky et al., 2017). These molecular differences alter the extracellular domains available for binding synaptic ligands and can change the effects of neurexins on synapse organization and transmission (Südhof, 2017). Individual neuronal cell types express reproducible combinations of neurexin genes, promoter-derived isoforms, and alternatively spliced variants, suggesting that neurexin expression is regulated as a component of neuronal identity (Fuccillo et al., 2015). These profiles can be established early in development and maintained during neuronal maturation (Lukacsovich et al., 2019). Experimental alteration of cell-type-specific neurexin splicing can disrupt synaptic and circuit function (Nguyen et al., 2016). Neurexins are further diversified by variable post-translational modification with heparan sulfate, which contributes to ligand binding and synaptic organization (Zhang et al., 2018). Recent work demonstrates that the minimal mammalian Nrxn1γ isoform uses its extracellular heparan sulfate glycans to bind AMPA receptors and align postsynaptic receptor nanodomains with presynaptic release sites (Padmanabhan et al., 2026). Together, these studies suggest that the specific neurexin isoforms expressed by a neuron may contribute to its synaptic properties. However, determining how this diversity is deployed across individual neurons in an intact nervous system remains challenging.

The nematode *Caenorhabditis elegans* provides an opportunity to examine neurexin isoform expression across individually identifiable neuronal cell types. In contrast to vertebrates, *C. elegans* contains a single classical neurexin gene, *nrx-1*, which produces multiple transcripts encoding long NRX-1α and short NRX-1γ isoforms (Haklai-Topper et al., 2011; Kurshan et al., 2018). A reporter containing regulatory sequences from the *nrx-1* locus was previously found to be expressed broadly throughout the nervous system, but this reporter did not distinguish among the different transcripts produced by the locus (Haklai-Topper et al., 2011). Functional studies have demonstrated that these transcripts do not act interchangeably. The short γ isoform is required for active zone assembly, synaptic vesicle recruitment, and calcium-channel clustering at presynaptic sites (Kurshan et al., 2018), and can stabilize nascent presynaptic assemblies following the recruitment of intracellular active zone scaffolds (Frankel et al., 2025). Long α and short γ isoforms also make distinct contributions to the initiation and maintenance of behavioral responses to food deprivation (Bastien et al., 2023), and to chemosensory avoidance across developmental stages (Muirhead et al., 2024). Despite these functional differences, the cellular expression patterns of the distinct *nrx-1* transcript groups have not been systematically resolved. It therefore remains unclear whether the broad expression of *nrx-1* reflects uniform coexpression of its transcripts or whether individual neuronal cell types deploy distinct subsets of *nrx-1* isoforms.

Here, we combined endogenous isoform-selective mutations with transcriptional reporters to examine how long α and short γ NRX-1 isoforms contribute to synaptic localization and to generate a nervous system-wide atlas of nrx-1 isoform expression. We find that both isoform classes contribute to NRX-1 protein localization at dorsal nerve cord synapses, with the short isoforms making the predominant contribution. Reporters corresponding to three groups of nrx-1 transcripts reveal distinct cellular and developmental expression patterns. Overall, the short γ isoforms are expressed more broadly, whereas the long α isoforms are expressed more selectively in particular neurons and anatomical regions. The two γ transcript groups also show distinct, partially overlapping expression patterns. We find striking differences in reporter expression between cholinergic and GABAergic motor neurons in the ventral nerve cord, as well as potential left-right differences in paired neurons. Finally, we identify distinct cis-regulatory elements that contribute to nrx-1m expression in different neuronal populations.

Together, our findings show that the broad neuronal expression of nrx-1 encompasses distinct combinations of long and short transcript groups in different neurons. These results map the cellular distribution of nrx-1 transcript groups and identify regulatory sequences contributing to their expression, providing a basis for testing how isoform expression contributes to the diverse synaptic functions of neurexin.

## RESULTS

### γ/Short NRX-1 isoforms make the predominant contribution to synaptic NRX-1 in the dorsal nerve cord

The *C. elegans nrx-1* locus has 13 annotated protein-coding transcripts, which are predicted to encode long NRX-1α and short NRX-1γ isoforms (Supplementary Fig. 1). To determine how these isoform classes contribute to synaptic NRX-1 localization, we generated targeted isoform deletions in the constitutive endogenously tagged *nrx-1::Skylan-S* allele. Skylan-S was previously inserted within the shared intracellular domain of all isoforms, immediately before the C-terminal PDZ binding motif, allowing all NRX-1 isoforms to be visualized (Kurshan et al., 2018). The *nrx-1::Skylan(Δα)* allele selectively removes the α isoforms while preserving the γ isoforms, whereas the *nrx-1::Skylan(Δγ)* allele is predicted to disrupt eight of the nine annotated γ transcripts, while preserving the α isoforms (Figure 1A).

**Figure 1.**
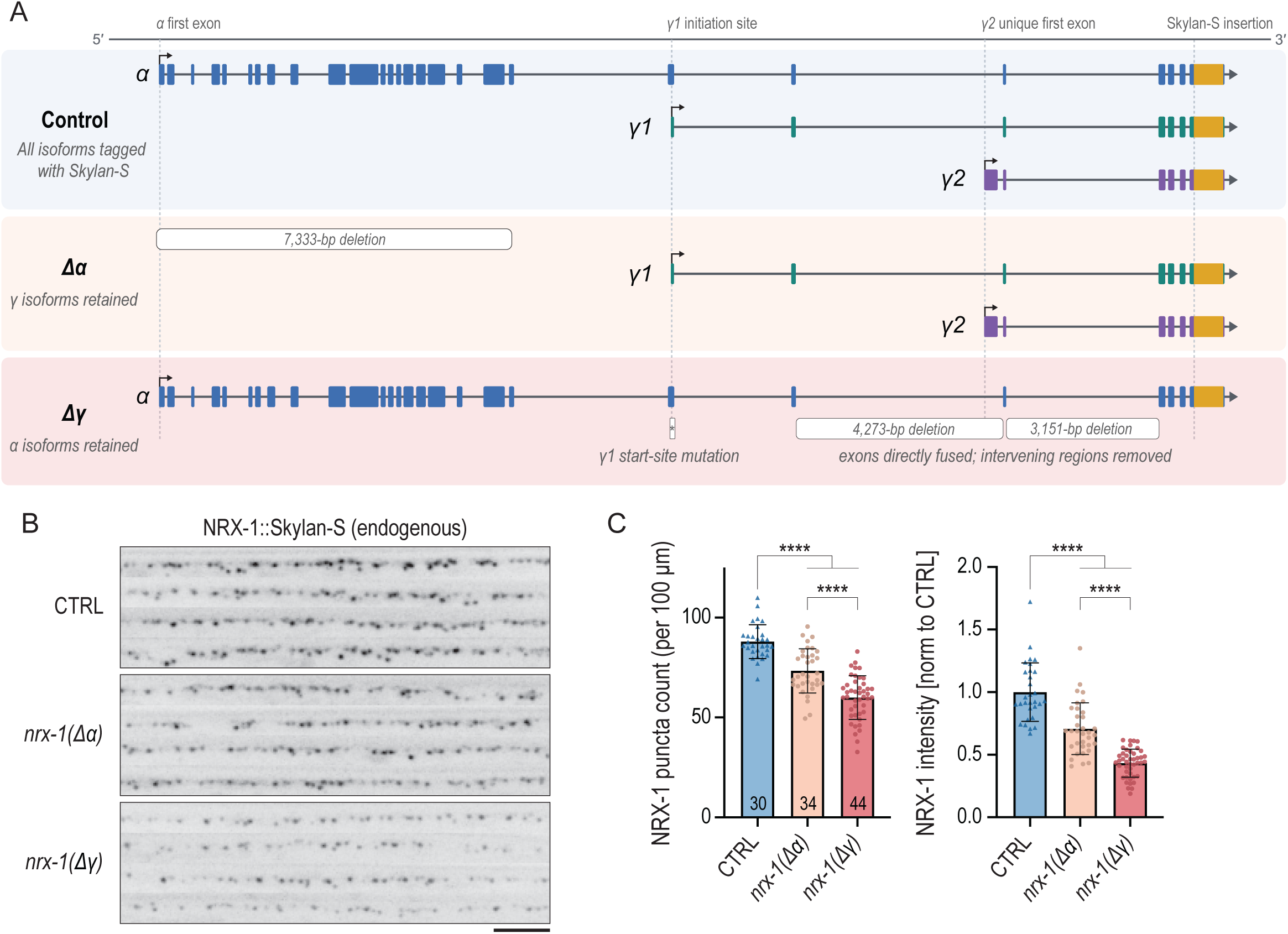
Short NRX-1 isoforms make the predominant contribution to synaptic NRX-1 in the dorsal nerve cord. **(A)** Schematic of the endogenous *nrx-1* locus and the isoform-selective alleles used to assess the contributions of long (*α*) and short (*γ*) isoforms to synaptic NRX-1 localization. Representative α, γ1, and γ2 transcripts and the Skylan-S insertion site are shown. **(B)** Representative maximum-projection images of endogenously labeled NRX-1::Skylan-S at dorsal nerve cord synapses in L4 hermaphrodites. CTRL animals contain all NRX-1 isoforms; *nrx-1(Δα)* animals retain only the short isoforms; and the *nrx-1(Δγ)* allele removes eight of the nine annotated short isoforms. Scale, 5 μm. **(C)** Quantification of NRX-1::Skylan synaptic puncta number and total punctal fluorescence per 100 μm of dorsal nerve cord. Fluorescence values were normalized to the CTRL mean. Numbers at the base of the puncta-count bars indicate the number of animals analyzed, and each point represents an individual animal. Bars indicate mean ± SD. Statistical significance was determined by Brown-Forsythe and Welch ANOVA followed by Dunnett’s T3 multiple comparisons test. Comparisons with CTRL are indicated; ****p<0.0001.

In L4 hermaphrodites, disruption of either isoform class reduced the density and total fluorescence of NRX-1 synaptic puncta in the dorsal nerve cord (Fig. 1B,C). Control animals exhibited approximately 87 NRX-1 puncta per 100 μm. Consistent with our previous analysis of *nrx-1::Skylan(Δα)* animals (Frankel et al., 2025), deletion of the α isoforms reduced puncta density to approximately 73 puncta per 100 μm and total punctal fluorescence to ∼71% of control. Disruption of the γ isoforms produced a larger reduction, to approximately 59 puncta per 100 μm and 43% of control fluorescence (Fig. 1C). Thus, both isoform classes contribute to synaptic NRX-1 localization, with the short γ isoforms making the predominant contribution to NRX-1 puncta density and fluorescence at dorsal nerve cord synapses.

### Isoform-specific reporters reveal the cellular distribution of NRX-1 expression

Short γ NRX-1 isoforms make the predominant contribution to NRX-1 localization at dorsal nerve cord synapses, but whether the α and γ isoforms are expressed in the same neurons or exhibit distinct expression patterns across the nervous system remained unclear. To define the cellular expression associated with different groups of *nrx-1* transcripts, we generated three endogenous GFP::T2A reporters. The α reporter marks *nrx-1a, -1b, -1k,* and *-1l*, the γ1 reporter marks *nrx-1c* and *-1d*, and the γ2 reporter marks *nrx-1f, -1g, -1i, -1j,* and *-1m* (Supplementary Fig. 1).

To generate an atlas of *nrx-1* isoform expression across the nervous system, we crossed our transcriptional reporters into the NeuroPAL background, allowing us to identify individual neurons by their fluorescence color and anatomical position (Fig. 2A). We compiled observations from L4 hermaphrodites for 283 neuron identities. For each neuron and reporter, we summarized detection as consistent across scored observations, variable across observations, or undetected. Combinations without a usable observation were marked as unassessed (Fig. 2E–G, Supplementary Fig. 2). Observation counts varied among neuron–reporter combinations and are provided in Supplementary Table 1. Among the 267 neurons assessed for all three reporters, 172 showed detection of multiple reporters, 83 showed detection of one, and 12 showed no detection of any reporter (Fig. 2B). The γ reporters were collectively detected more broadly than α, and most neurons with α detection also showed detection of at least one γ reporter (Fig. 2C). The two γ reporters also differed in their distributions, with both detected in 90 neurons, γ1 without γ2 in 77, and γ2 without γ1 in 76 (Fig. 2D).

**Figure 2.**
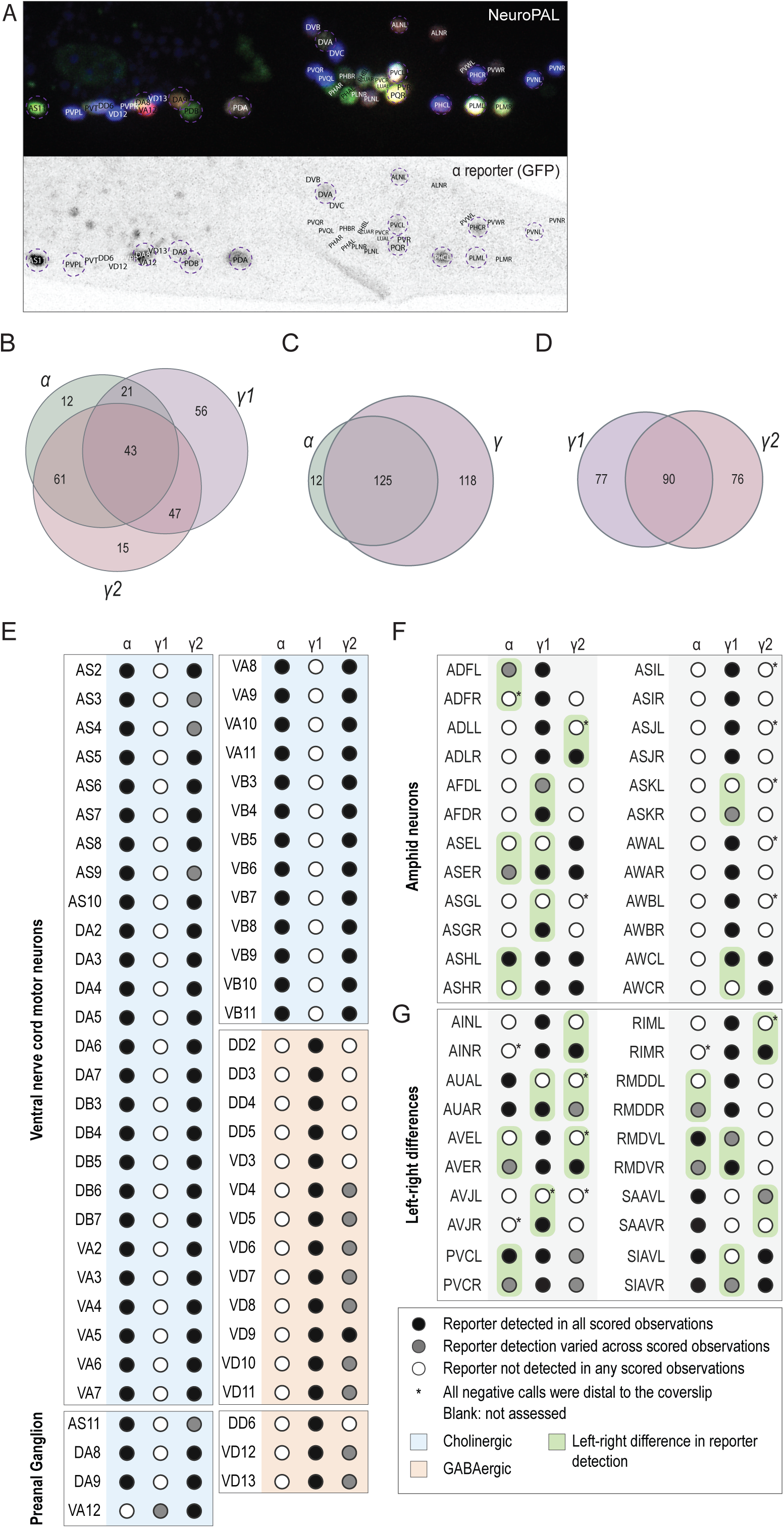
NeuroPAL-based map of *nrx-1* isoform-group reporter expression in individually identified neurons. Reporter detection is summarized for the α (*nrx-1a/b/k/l*), γ1 (*nrx-1c/d*), and γ2 (*nrx-1f/g/i/j/m*) groups. **(A)** NeuroPAL fluorescence (top) and corresponding α-reporter GFP signal (bottom), with identified posterior neurons labeled. **(B–D)** Overlap of reporter detection among 267 neurons assessed for all three reporter groups: α, γ1, and γ2 individually (B); α versus either γ reporter (C); and γ1 versus γ2 (D). Detection in at least one scored observation qualifies a neuron for inclusion in a reporter set. Numbers indicate neuron counts. Neurons without detection in any displayed set are omitted: 12 neurons in B and C, and 24 in D. **(E)** Ventral nerve cord and preanal ganglion motor neurons, with cholinergic and GABAergic identities color-coded blue and peach, respectively. **(F,G)** Reporter detection in amphid neurons (F) and selected additional neuron pairs (G). Green highlights indicate potential left-right asymmetries in reporter detection, including comparisons involving variable detection. In panels E-G, each circle summarizes reporter detection across scored L4 hermaphrodites. Black circles indicate detection in every scored observation; gray circles indicate both positive and negative observations; and open circles indicate no detection in any scored observation. Blank entries indicate that no usable detection calls were available and do not establish lack of expression. Asterisks identify assignments for which all recorded negative observations were obtained with the neuron on the side farther from the coverslip. Sample sizes varied among neuron-reporter combinations; observation counts are provided in Supplementary Table 1. Additional neurons and numerical fluorescence measurements for a subset of neurons are shown in Supplementary Figures 2 and 3, respectively.

The clearest class-associated pattern emerged in the ventral nerve cord motor neurons (Fig. 2E). The α reporter was detected in all 39 cholinergic motor neurons examined and was undetected in all 13 GABAergic motor neurons. Conversely, γ1 was detected in all 13 GABAergic neurons and was undetected in all 39 cholinergic neurons. The γ2 reporter was consistently detected across observations in 36 cholinergic neurons and variably detected in the remaining three. Among the GABAergic neurons, γ2 was consistently detected in one, variably detected in seven, and undetected in five. Most preanal ganglion motor neurons followed the same α/γ1 distinction. VA12 was an exception, with α undetected and γ1 variably detected. Together, these observations reveal distinct *nrx-1* reporter profiles in cholinergic and GABAergic motor neurons.

The association between neurotransmitter identity and reporter expression did not extend uniformly beyond the ventral nerve cord. For example, γ1 was detected in the cholinergic IL2 neurons and in the glutamatergic ASER neuron (Fig. 2F, Supplementary Fig. 2). Dopaminergic neurons also showed different profiles. The γ2 reporter was detected in CEPDL and CEPDR, where α was undetected, whereas PDEL and PDER showed detection of all three reporters. Thus, neurotransmitter identity alone did not predict a uniform reporter profile across the nervous system.

Our atlas also revealed regional differences. The α reporter was undetected in all six dorsal ganglion neurons examined, although several showed γ1 or γ2 detection. Amphid sensory neurons predominantly showed detection of the short γ reporters, whereas all three reporters were detected in each of the 12 assessed midbody neurons (Fig. 2F, Supplementary Fig. 2). Several bilateral pairs differed in reporter detection between their left and right members (Fig. 2F,G). For example, γ1 was detected in ASER and AWCL but was undetected in ASEL and AWCR. Other pairs differed in the consistency of detection across animals. Across the 267 neurons assessed for all reporters, 12 showed no detection of any reporter. These neurons may express *nrx-1* below the detection limit or express transcripts outside the groups captured by these reporters.

For a subset of neurons, software-derived GFP estimates also allowed us to examine fluorescence intensity within each reporter line (Supplementary Fig. 3, Supplementary Table 1). These measurements included midbody and posterior neurons, as well as ASE neurons. Among posterior neurons, γ2 fluorescence was relatively high in VA12, PDA, and DA8, with lower estimates in PVQ and PVW neurons. The α and γ1 reporters also showed variation among neurons. These measurements extend the detection map by describing variation in reporter fluorescence within each reporter group.

To compare the reporter patterns with neuronal RNA-seq data, we examined CeNGEN coverage near the isoform starts corresponding to the three reporter groups in 51 matched neuron classes (Supplementary Fig. 4, Weinreb et al., 2025). We compared relative RNA signals within each class with the fraction of L4 observations in which each reporter was detected. The DA and DD motor neurons illustrated correspondence between the datasets. DA showed its highest relative RNA signal for α and detection of the α and γ2 reporters, whereas DD showed its highest relative RNA signal for γ1, the only reporter detected in that class. Across all 51 classes, correlations between reporter detection frequency and relative RNA signal were modest for α and γ1 and weaker for γ2 (Spearman’s ρ = 0.43, 0.33, and 0.23, respectively).

### *The α and γ2 nrx-1 reporters* exhibit distinct developmental and cellular expression patterns

To compare isoform expression patterns directly within the same animal, we generated a dual reporter combining the α GFP::T2A reporter with an mScarlet::SL2 reporter upstream of *nrx-1m.* In the heads of L4 hermaphrodites, the two reporters were co-expressed in several cell bodies, but each reporter was also detected in cells with little or no detectable expression of the other reporter (Fig. 3A). These results indicate that the α and γ2 reporters are expressed in partially overlapping but distinct populations of cells and suggest that the two isoforms are subject to distinct transcriptional regulation.

**Figure 3.**
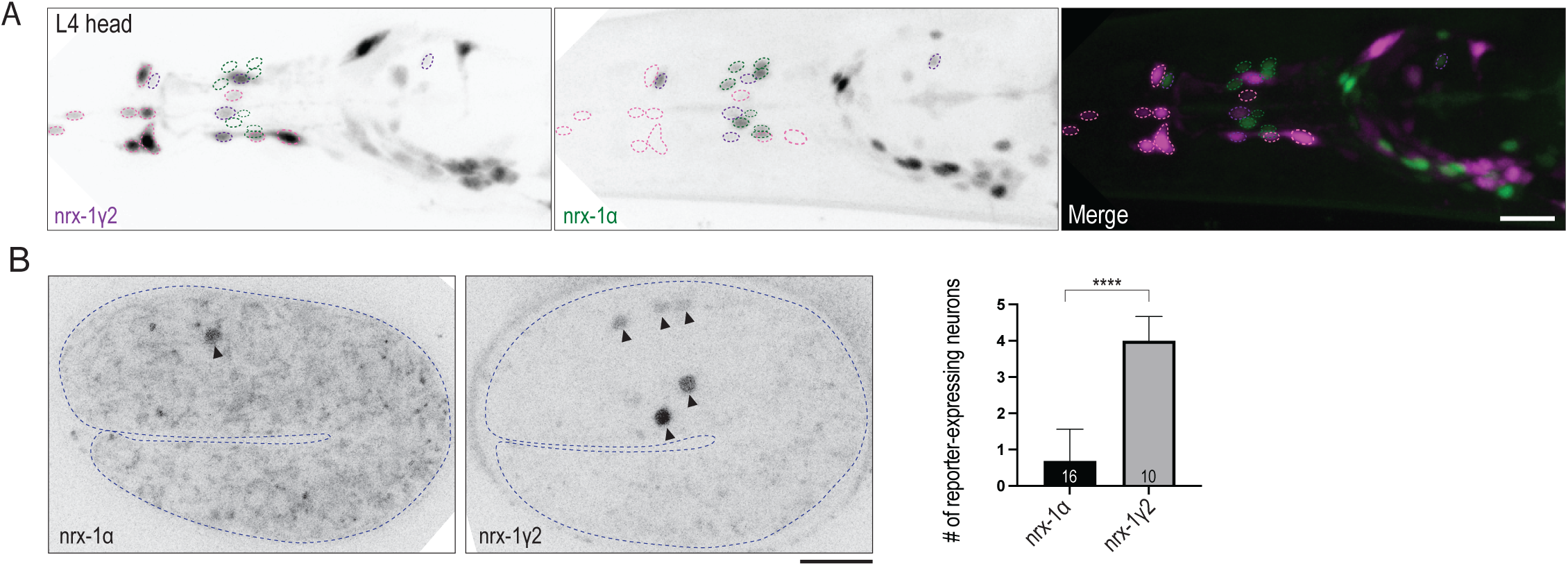
*nrx-1α* and *nrx-1γ2* exhibit distinct developmental and cellular expression patterns. **(A)** Representative maximum-projection images of the head of an L4 hermaphrodite carrying the *nrx-1a* GFP::T2A reporter (α) and the *nrx-1m* mScarlet::SL2 (γ2). α and γ2 reporter signals are pseudocolored green and magenta, respectively, in the merged image. Colored outlines highlight representative cell bodies with predominantly *nrx-1α* reporter expression (green), predominantly *nrx-1γ2* reporter expression (pink), or detectable expression of both reporters (purple). Scale bar, 10 μm. **(B)** Representative maximum-projection images of α (*nrx-1a)* and γ2 (*nrx-1m*) GFP::T2A reporter expression in two-fold embryos, displayed in inverted grayscale. Arrowheads indicate reporter-expressing neuronal cell bodies. The graph shows the number of GFP-positive neuronal cell bodies per embryo. Numbers at the base of each bar indicate the number of embryos analyzed. Bars indicate mean ± SD. Groups were compared using an unpaired, two-tailed Welch’s *t-*test (****P<0.0001). Scale bar, 10 μm.

To examine developmental differences in isoform expression, we compared the α and γ2 GFP::T2A reporters in two-fold embryos. The α reporter was detected in relatively few neuronal cell bodies at this stage, with an average of 0.68 reporter-expressing neurons per embryo. In contrast, the γ2 reporter was detected in approximately four neuronal cell bodies per embryo (Fig. 3B). Thus, γ2 reporter expression was more widespread than α reporter expression by the two-fold stage of embryogenesis.

### Distinct cis-regulatory elements control *nrx-1m* expression in different neuronal populations

Because the γ2 reporter exhibited a cellular expression pattern partially distinct from that of α, we next investigated the cis-regulatory sequences contributing to *nrx-1m* expression. UNC-3, the *C. elegans* COE-family terminal selector, broadly regulates cholinergic motor neuron differentiation and was previously shown to control *nrx-1α* expression through a COE motif (Philbrook et al., 2018). We therefore tested whether a predicted COE motif upstream of *nrx-1 (γ2)* contributes to reporter expression. We also deleted an approximately 200-bp upstream sequence containing predicted binding sites or overlapping ChIP-Seq peaks for LIN-14, EGL-13, UNC-62, FOS-1, and NHR-6, hereafter referred to as the γ2 upstream regulatory element (γ2URE, Fig. 4A)

**Figure 4.**
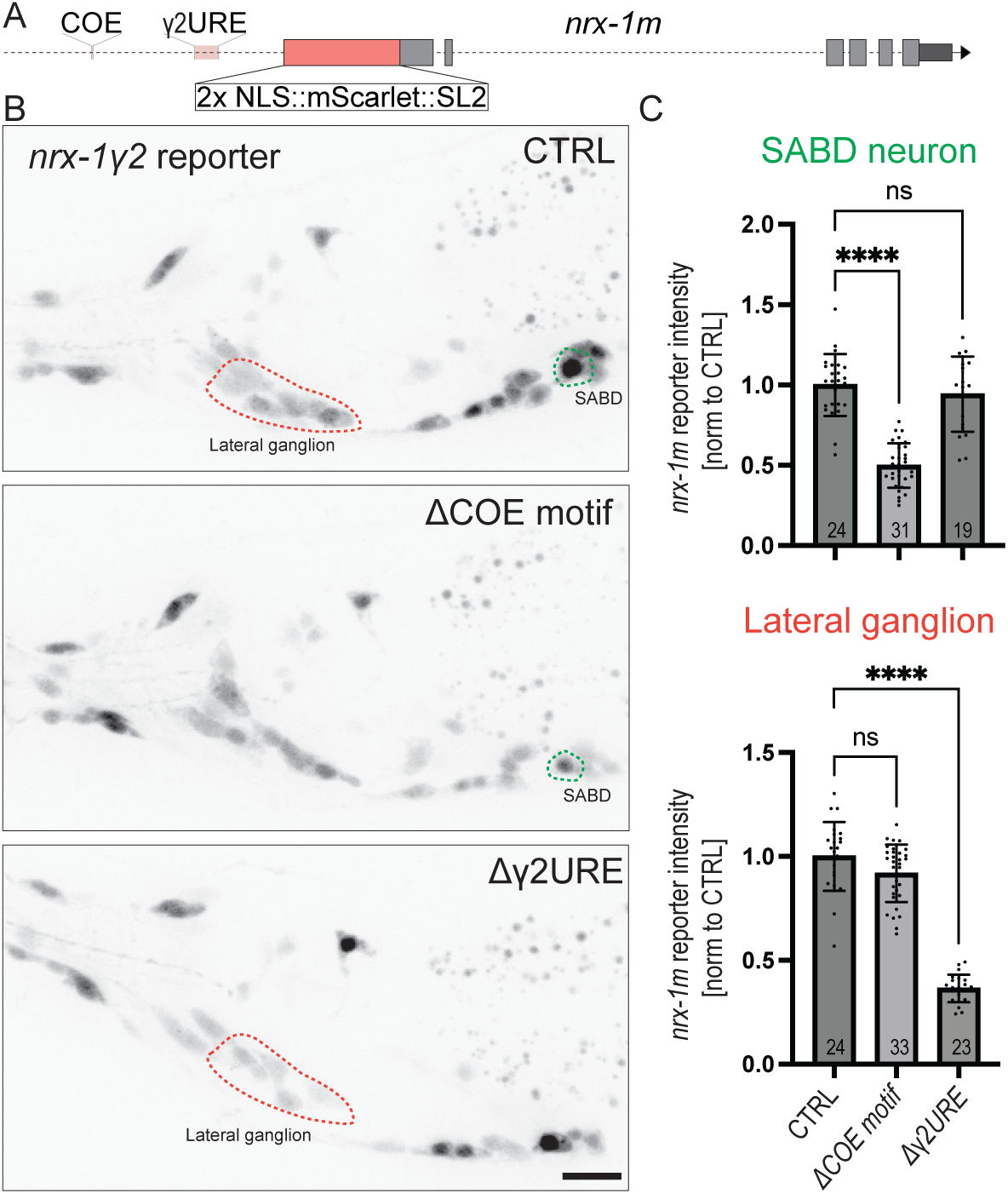
Distinct cis-regulatory elements contribute to *nrx-1m* expression in different neuronal populations. **(A)** Schematic of the endogenous *nrx-1m* transcriptional reporter, containing a 2xNLS::mScarlet::SL2 cassette, and the positions of upstream COE motif and γ2 regulatory element (γ2URE) targeted for deletion. **(B)** Representative maximum-projection images of γ2 reporter expression in the heads of L4 hermaphrodites carrying the control reporter allele (CTRL), the COE motif deletion (ΔCOE), or the γ2URE deletion (Δγ2URE). Green and red outlines mark the neuron tentatively identified as SABD and the lateral ganglion region used for fluorescence quantification, respectively. Scale bar, 10 μm. **(C)** Quantification of γ2 reporter fluorescence in putative SABD and the lateral ganglion. Fluorescence values were normalized to the CTRL mean for each neuronal population. Numbers at the base of each bar indicate the number of animals analyzed, and each point represents an individual animal. Bars indicate mean ± SD. Statistical significance was determined by Brown-Forsythe and Welch ANOVA followed by Dunnett’s T3 multiple comparisons test. Comparisons with CTRL are indicated; ns, not significant; ****p<0.0001.

The two regulatory sequences affected *nrx-1(γ2)* reporter expression in different neuronal populations. Deletion of the COE motif reduced reporter fluorescence in a neuron tentatively identified as SABD to approximately 50% of control but did not significantly affect expression in the lateral ganglion (Figure 4B,C). Because SABD is a cholinergic motor neuron, this selective effect is consistent with UNC-3 contributing to *nrx-1(γ2)* expression through the COE motif.

Conversely, deletion of the γ2URE did not significantly affect expression in putative SABD but reduced aggregate reporter fluorescence in the lateral ganglion to approximately 36% of control. Thus, distinct cis-regulatory elements contribute to *nrx-1 γ2* expression in different neuronal populations.

## DISCUSSION

Neurexins are synaptic cell-adhesion molecules with central roles in synapse development, organization, and function, and their disruption is associated with neurodevelopmental disorders, including autism and schizophrenia. Neurexins are expressed as isoforms of different lengths. Although the α and β isoforms have been extensively characterized with respect to their functions and binding partners, much less is known about the shorter γ isoforms. Here, we generated a nervous system-wide atlas of neurexin isoform expression and identified distinct expression patterns for each isoform class. We found that the short neurexin isoforms are collectively expressed more broadly throughout the *C. elegans* nervous system, that the γ2 reporter is expressed in more neurons than α at the two-fold embryonic stage, and that separable cis-regulatory elements contribute to γ2 expression in distinct neuronal populations. To our knowledge, this work represents the first nervous-system-wide atlas of isoform-specific expression for this important synaptic gene.

### An atlas of long and short NRX-1 isoform expression reveals distinct patterns

Our atlas reveals broad expression of the short γ transcript groups, alongside more selective expression of the long α group. Among the 267 neurons assessed for all three reporters, 91% showed detection of at least one γ reporter, compared with 51% for α. Most neurons with α detection also showed detection of a γ reporter. This distribution suggests that the short isoforms provide functions shared across many neuronal populations, while the addition of α supports further specialization.

We previously showed that, in *C. elegans*, short NRX-1γ can stabilize nascent presynaptic assemblies in the dorsal nerve cord (Frankel et al., 2025). We therefore proposed that this isoform may provide a minimal neurexin function required for presynaptic stabilization before transsynaptic alignment or the establishment of postsynaptic partner specificity. Here, we find that γ isoforms are broadly expressed across the L4 nervous system, and that the γ2 reporter expression is more widespread than α at the two-fold embryonic stage. These findings support a model in which short γ isoforms provide a broadly deployed foundation for stabilization of nascent presynaptic assemblies, with their effects shaped by additional neurexin isoforms, cell-specific ligands, and postsynaptic context. In this model, selective expression of the α isoform would add synapse-specific functions to a more broadly shared γ-dependent program. Whether γ neurexin performs a common function across different circuits remains to be determined.

Although the two γ transcript groups encode similarly short neurexin isoforms, their expression patterns differed. The γ1 reporter was detected in most amphid sensory neurons, whereas γ2 was restricted to a smaller subset. Ventral nerve cord motor neurons were split by neurotransmitter identity, while assessed midbody neurons frequently expressed both short isoforms alongside α. These patterns suggest that neuronal identity influences both the expression of long versus short neurexins and the deployment of distinct short transcript groups.

The ventral nerve cord motor neurons provide a particularly striking example of isoform-specific expression. The long α isoform is expressed by cholinergic but not GABAergic motor neurons, whereas the γ1 isoform exhibits the reciprocal pattern. Both motor-neuron types innervate body-wall muscle, and cholinergic motor neurons also synapse onto GABAergic motor-neuron dendrites. Previous work showed that long NRX-1α regulates acetylcholine receptor clustering on GABAergic motor-neuron dendrites, but not on body-wall muscles (Philbrook et al., 2018). The selective expression of α in cholinergic motor neurons is therefore consistent with a role in organizing their synapses onto GABAergic neurons.

Distinct isoform-expression patterns are also evident among sensory neurons. Amphid sensory neurons, which include chemosensory, olfactory, thermosensory, and polymodal neurons, predominantly express γ isoforms, whereas midbody mechanosensory neurons robustly express all three isoforms. Anatomical position may also contribute to these differences. For example, α and γ2 were consistently detected in anterior ALM neurons but variably detected in posterior PLM neurons. Among posterior motor neurons, VA12 departed from the VNC α/γ1 pattern, with α undetected and γ1 variably detected. These observations suggest that isoform expression may reflect an intersection of functional, neuronal-class, and positional identity. They further raise the possibility that regional patterning programs, including Hox-dependent mechanisms (Kratsios et al., 2017; Zheng et al., 2022, 2015), modify cell-type-specific regulation of the *nrx-1* locus.

Our atlas also identified potential left-right asymmetries in neurexin isoform expression, involving both long and short transcript groups. For example, γ1 was detected in ASER and AWCL but was undetected in ASEL and AWCR. Among head-neuron pairs with different isoform expression, the right member more often showed detection or more consistent detection of the γ reporters. Unequal imaging orientations could contribute to this apparent bias, particularly where negative observations were obtained from neurons farther from the coverslip. However, left–right differences in isoform expression may reflect lateralized requirements associated with synaptic partner specificity or strength. For example, α was variably detected in ADFL and was undetected in ADFR. These neurons also differ in their patterns of postsynaptic connectivity. The adult hermaphrodite connectome includes ADFL outputs onto AUAL and RIBL and ADFR outputs onto RIGR and URXR (Cook et al., 2019). Whether differences in α expression contribute to the synaptic organization of the ADF pair remains unknown.

### Quantitative analysis and comparison to CeNGEN RNA-seq data

Our fluorescence measurements further suggest that variation in *nrx-1* expression extends beyond differences in reporter detection. Estimated GFP levels varied among neurons within each reporter group, including neurons in which the reporter was consistently detected (Supplementary Fig. 3). Thus, neuronal differences in neurexin expression may involve both the combination of transcript groups expressed and their relative transcriptional output. Whether differences in reporter intensity correspond to differences in endogenous transcript abundance, synaptic protein abundance, or function remains to be determined.

Comparison with CeNGEN short-read RNA-seq data revealed partial correspondence with the reporter patterns (Hammarlund et al., 2018; Weinreb et al., 2025). The distinction between DA and DD motor neurons was evident in both datasets, but agreement across the 51 matched neuron classes was modest for α and γ1 and weaker for γ2 (Supplementary Fig. 4). One challenge is that coverage near an internal isoform start can include reads originating from longer transcripts. Our analysis therefore used coverage at the α coding start, α-subtracted coverage at the γ1 coding start, and the local rise in coverage at the γ2 transcript start to estimate relative signals. These estimates were compared with reporter detection frequencies pooled within the corresponding neuron classes. Differences between the datasets could reflect limitations in assigning shared reads to transcript groups, differences in detection sensitivity, or biological variation within a neuron class. The comparison identifies neuronal populations with convergent evidence for particular expression patterns and others where direct transcript measurements would help resolve the discrepancies.

### Transcriptional regulation of neurexin isoforms

Our cis-regulatory analysis suggests that distinct regulatory elements contribute to *nrx-1m* expression in different neuronal populations. Deleting the predicted COE motif reduced reporter fluorescence in putative SABD, while lateral-ganglion reporter fluorescence did not differ significantly from control animals. Deleting the γ2URE reduced lateral-ganglion reporter fluorescence, with no significant effect in putative SABD. These findings support a model in which modular regulation at the *nrx-1m* promoter generates expression across different neuronal populations and adjusts expression levels within them. More generally, the combinatorial action of regulatory elements at individual promoters may contribute to the cellular diversity of neurexin expression alongside alternative promoter use and splicing.

### Caveats and open questions about neurexin isoform expression

Our atlas maps the expression of groups of nrx-1 transcripts but does not resolve individual transcripts within each group. Reporter fluorescence provides an indication of transcriptional activity, while the abundance and localization of the encoded NRX-1 proteins remain to be established. Isoform-specific protein measurements will therefore be important for determining how the expression patterns identified here translate into differences at synapses.

An additional question is whether isoforms expressed within the same neuron localize to distinct presynaptic specializations contacting different postsynaptic partners. Our dual-reporter experiment demonstrates simultaneous α and γ2 reporter expression in some L4 head neurons. However, the nervous system-wide map combines observations from separate reporter strains, leaving open how frequently all three transcript groups are co-expressed within individual animals. Longitudinal imaging with spectrally distinct reporters could also establish whether neurons switch between transcript groups or progressively add them during development. Combining these approaches with isoform-specific protein tags would connect changes in expression with the distribution of NRX-1 at individual synapses.

The fluorescence reductions caused by the two isoform-selective mutations did not sum to the total control signal. The Δγ allele is predicted to disrupt eight of the nine annotated short transcripts, so residual expression from the remaining transcript could contribute to this difference. The mutations could also induce compensatory changes in expression or reveal interactions between α and γ isoforms that affect trafficking or synaptic retention. Isoform-specific protein measurements in the mutant backgrounds would help distinguish among these possibilities.

Our data also leave open the circuit and behavioral consequences of losing individual neurexin isoforms. This could be assessed by examining behaviors controlled by asymmetric neuronal pairs that differ in reporter expression. For example, the ASE salt-sensing circuit undergoes asymmetric changes in synaptic connectivity during associative learning (Tang et al., 2023). We detected γ1 in ASER and not in ASEL, raising the possibility that differences in isoform expression contribute to functional or plasticity differences within this circuit. Testing that possibility will require selective manipulation of the transcript groups in each neuron.

Overall, our findings establish a cellular map of long and short nrx-1 transcript-group expression and identify regulatory elements that contribute to expression in distinct neuronal populations. This atlas provides a resource for selecting neurons and circuits in which to test how particular isoform combinations contribute to synapse development, maintenance, and plasticity.

## METHODS

### *C. elegans* strains and maintenance

*C. elegans* strains were maintained at 20°C or 23°C on NGM plates seeded with *E. coli* OP50 using standard procedures (Brenner, 1974). Experiments were performed using hermaphrodites at the developmental stages specified below.

### Generation of CRISPR-Cas9 alleles

Endogenous *nrx-1* alleles were generated with gonadal injection of CRISPR-Cas9 ribonucleoprotein complexes as previously described (Frankel et al., 2025) . Injection mixes contained 1.525 μM Alt-R Cas9-tracrRNA-crRNA complexes (Integrated DNA Technologies) and either 6 μM single-stranded DNA repair templates or 1-5 μM PCR-amplified repair templates, depending on the size and complexity of the edit. A co-CRISPR strategy targeting *dpy-10* or *unc-58* was used to facilitate identification of edited animals(Arribere et al., 2014). Repair templates contained 100-200 bases of homology on each side of the targeted region. Where applicable, silent substitutions were introduced into the protospacer adjacent motif (PAM) or guide RNA recognition sequence to prevent recutting. Edited alleles were identified by PCR screening, verified by Sanger sequencing.

### Fluorescence microscopy

Fluorescence images, including those used for NeuroPAL neuronal identification, were acquired using a spinning-disk confocal system (3i) comprising an inverted Zeiss Axio Observer Z1 microscope, a Yokogawa CSU-X1 M1 spinning disk, and a Photometrics Evolve 512 EMCCD camera.

For dorsal nerve cord and embryonic imaging, images were acquired at room temperature using a Zeiss Plan-Apochromat 100×/1.4 NA oil-immersion objective and SlideBook 6.0 software (3i). L4 hermaphrodites were immobilized in 20 mM levamisole in M9 buffer, mounted on 10% agarose pads, and sealed beneath coverslips with petroleum jelly. Animals were staged by vulval morphology using differential interference contrast (DIC) microscopy, and those between stages L4.4 and L4.6 were selected for imaging (Mok et al., 2015). Dorsal nerve cord z-stacks were acquired at 0.27 μm intervals over a 6 μm range.

Embryos were mounted on 10% agarose pads and staged according to morphology. Embryonic z-stacks were acquired at 0.3 μm intervals over a 6 μm range. For NeuroPAL imaging, L4 hermaphrodites carrying an endogenous nrx-1 GFP::T2A reporter and the NeuroPAL transgene otIs669 V were immobilized in 20 mM levamisole in M9 buffer and mounted on 10% agarose pads. Animals were positioned laterally or dorsoventrally as needed to resolve neurons in the regions examined. Multichannel confocal z-stacks encompassing neuronal cell bodies and surrounding NeuroPAL landmarks were collected using a 63×/1.45 NA objective at 1 μm intervals. Excitation wavelengths and emission filters were selected according to the published NeuroPAL protocol (Yemini et al., 2021).

### Image processing and fluorescence quantification

Dorsal nerve cord confocal z-stacks were converted to 16-bit maximum-intensity projections. Synaptic regions were traced in Fiji using 20-pixel-wide lines and straightened to generate 32-bit TIFF images for analysis (Schindelin et al., 2012). Fluorescent puncta were detected and segmented using WormSNAP, a custom MATLAB-based application(Tiroumalechetty et al., 2025). The same segmentation thresholds and filtering parameters were applied to all images of a given fluorescent marker across genotypes.

Puncta number and summed puncta fluorescence intensity were quantified and normalized to 100 μm of nerve cord. Reductions in detected puncta number may reflect both fewer puncta and puncta falling below the detection threshold. Representative images were selected to reflect the quantitative results for each genotype. For display, identical linear adjustments to brightness and contrast were applied to images compared across genotypes. Images were displayed using an inverted grayscale lookup table.

Two-fold embryos carrying the α or γ2 GFP::T2A reporter were identified by morphology and imaged as described above. GFP-positive neuronal cell bodies were counted manually from the acquired z-stack. A cell body was counted as positive when a discrete cell-body signal distinguishable from surrounding fluorescence was observed. Counts were recorded for individual embryos, with each embryo treated as a biological replicate.

Reporter fluorescence was quantified in putative SABD and the lateral ganglion of L4 hermaphrodites carrying the intact *nrx-1m* reporter, the COE-motif deletion, or the γ2URE deletion. Regions of interest were drawn around the putative SABD cell body and the lateral-ganglion region using FIJI. Mean fluorescence was measured from maximum intensity projections. No background subtraction was applied. For each neuronal population, measurements were normalized to the mean of control animals carrying the intact reporter allele. Each animal contributed one measurement per population, and animals were treated as biological replicates.

### Neuronal identification and reporter expression scoring

Neuronal identities were assigned using NeuroPAL fluorescence patterns and cell-body positions relative to neighboring neurons (Yemini et al., 2021). GFP expression from the endogenous nrx-1 α, γ1, and γ2 reporters was evaluated in identified neurons. Neurons that could not be confidently identified or assessed for GFP expression were treated as unscored rather than reporter-negative.

GFP expression was scored as present when fluorescence could be distinguished from local background and assigned to an identified neuron. Expression observations were compiled separately for each neuron and reporter group across animals. The number of animals scored for each neuron-reporter combination is provided in Supplementary Table 1.

Reporter brightness was initially assessed by visual inspection of fluorescence images and assigned a score from 0 to 3 (no GFP to bright GFP signal). Subsequent analyses used NeuroPAL Auto-ID, version 1 software to obtain estimated GFP values for identified neurons. The brightness summaries presented in Supplementary Table 1 were based on the automated software IDs. Comparisons were restricted to neurons within the same reporter group; brightness values are not comparable across the α, γ1, and γ2 reporters.

### Comparison with CeNGEN RNA-seq data

RNA-seq coverage data were obtained from the CeNGEN splicing portal (Weinreb et al., 2025), using the WS289 transcript annotation. All coordinates below refer to chromosome V, with the first base numbered 1 and both endpoints included. For α, mean coverage in counts per million (CPM) was calculated over the first 30 coding bases, spanning 10,825,024–10,825,053. For γ1, mean coverage was calculated over the first 30 coding bases of the *nrx-1c/d* transcripts, comprising 27 bases at 10,835,626–10,835,652 and three bases in the following exon at 10,838,067–10,838,069. These measurements used neuron-class mean coverage tracks. The α signal was used directly, and an adjusted γ1 signal was calculated by subtracting α coverage from γ1 coverage, with negative differences set to zero. For γ2, a local coverage change was calculated for each RNA-seq sample by subtracting mean coverage over the 10 bases immediately upstream of the annotated *nrx-1f* transcript start (10,842,022–10,842,031) from mean coverage over its first 10 bases (10,842,032–10,842,041). The median change across samples within each neuron class was used as the γ2 signal, with negative values set to zero.

The three RNA signals were divided by the largest signal within each neuron class, producing relative values from zero to one. Reporter observations were restricted to L4 hermaphrodites and grouped according to the corresponding CeNGEN neuron classes. For each class and reporter, detection frequency was calculated as the total number of positive observations divided by the total number of scored observations. Individual neurons, including bilateral partners and members of motor-neuron classes, were pooled where represented together in CeNGEN. The comparison included 51 classes with scored L4 observations for all three reporters. Spearman’s rank correlations were calculated across classes between each reporter’s detection frequency and its corresponding relative RNA signal.

### Quantification and statistical analysis

Quantitative data are presented as mean ± standard deviation (SD), with individual animal measurements overlaid, unless otherwise indicated. For synaptic measurements, each animal constituted one biological replicate, and sample sizes are indicated in the figures. Data were collected across multiple imaging sessions on multiple days, with control and experimental groups imaged together.

Statistical analyses were performed using GraphPad Prism (version 11.1). Pairwise comparisons were analyzed using unpaired t tests with Welch’s correction. Comparisons among multiple groups were analyzed using Brown–Forsythe and Welch ANOVA with Dunnett’s T3 multiple-comparisons tests, unless otherwise indicated. The statistical tests used for each analysis are specified in the corresponding figure legends. A p value < 0.05 was considered statistically significant.

NeuroPAL expression and software-derived GFP intensity data were summarized descriptively. Sample sizes represent the number of animals scored for each neuron–reporter combination and are provided in Supplementary Table 1. GFP intensity comparisons were restricted to neurons within the same reporter group.

Supplementary Table 1. Reporter detection, observation counts, and fluorescence intensities for *nrx-1* reporters in identified neurons. The table summarizes the α, γ1, and γ2 reporters in 283 neurons. “Scored n” gives the number of observations scored as detected or not detected, with positive and negative counts listed separately. Tentative calls (“y?” and “n?”) were included in their stated direction; their counts are subsets of the corresponding totals. Numerical fluorescence measurements were available for a subset of neurons and include measurements scored as reporter-negative. “Intensity n” gives the number of measurements used to calculate mean estimated GFP fluorescence, reported in arbitrary units. Blank mean-intensity cells indicate that no usable numerical measurements were available. All observations were from L4 hermaphrodites.

## ACKNOWLEDGEMENTS

We thank members of the Kurshan lab as well as Hannes Buelow, Scott Emmons and David Hall for helpful discussions. This work was funded by NINDS R01NS123645 (PTK) and the McKnight Foundation (PTK).

---

**Supplementary Figure 1.**
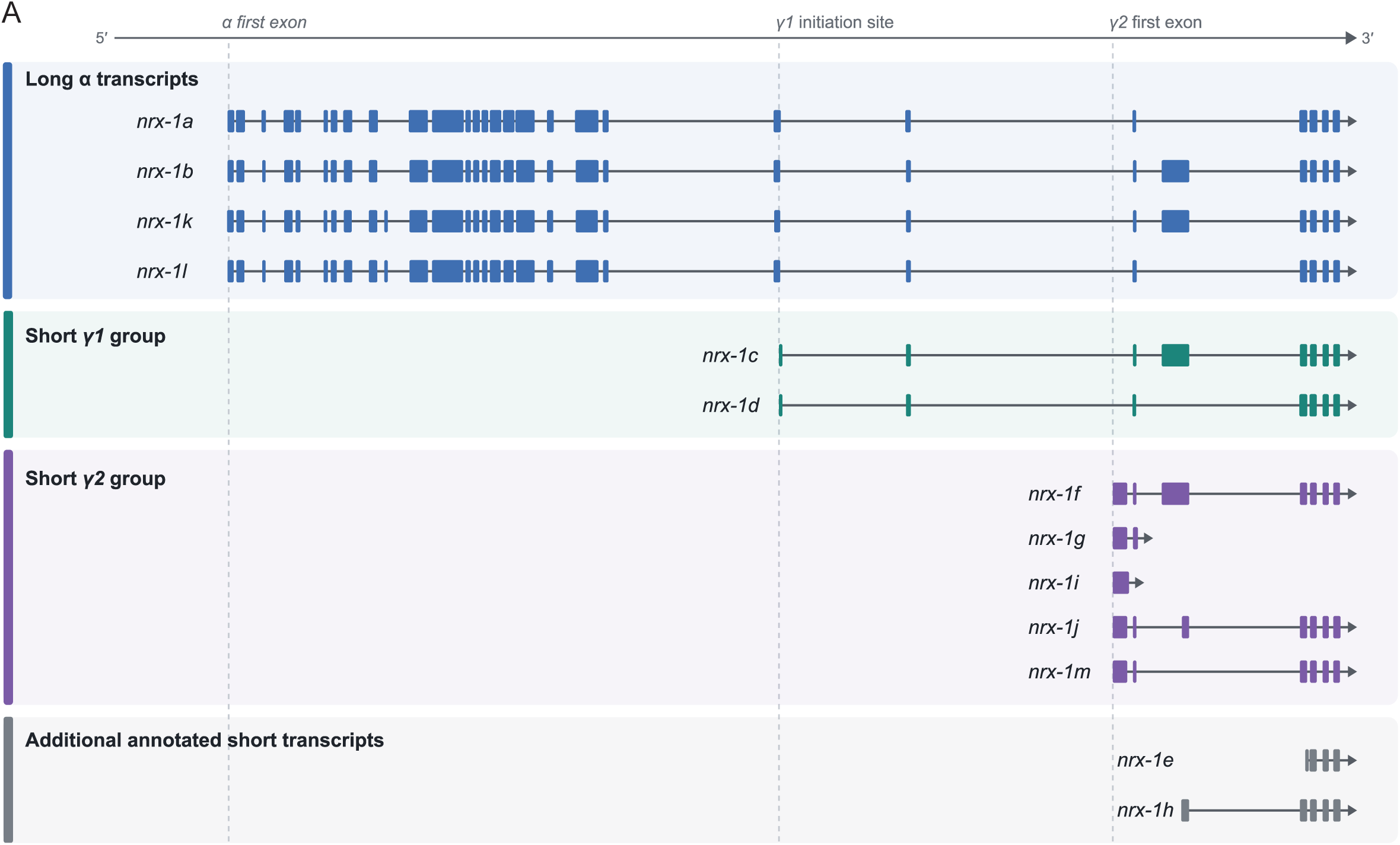
Organization of *nrx-1* transcript isoforms. Schematic of annotated *nrx-1* transcripts, grouped as long α transcripts (*nrx-1a, nrx-1b, nrx-1k, and nrx-1l), short* γ1 transcripts (*nrx-1c, nrx-1d)*, short γ2 transcripts (*nrx-1f, nrx-1g, nrx-1i, nrx-1j, nrx-1m*), and additional annotated short transcripts (*nrx-1e* and *nrx-1h*). Boxes indicate exons, connecting lines represent introns and arrows indicate the direction of transcription. Transcript groups are distinguished by color.

**Supplementary Figure 2.**
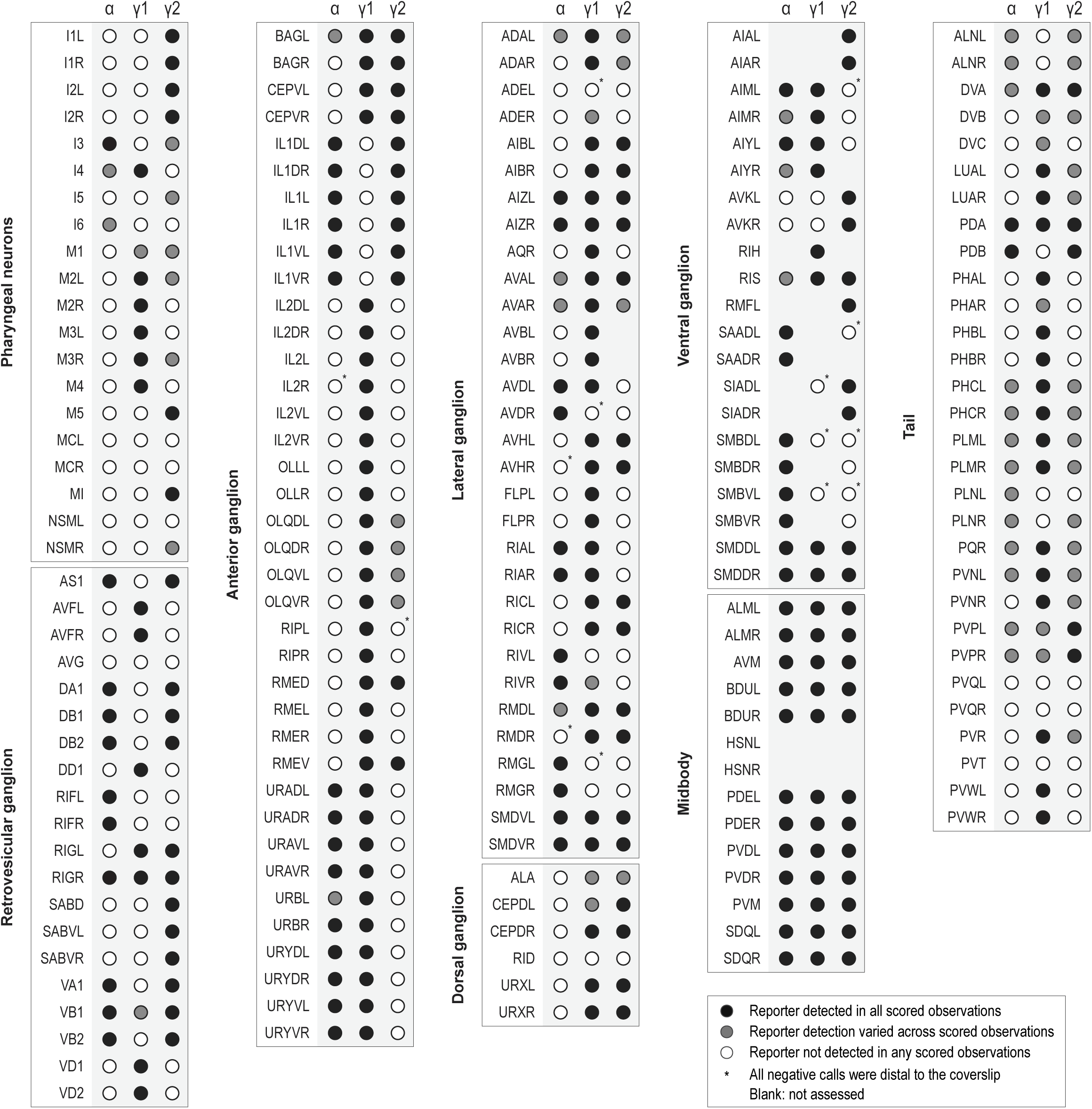
Detection of *nrx-1* reporters in neurons not shown in. Figure 2. Reporter detection was scored in L4 hermaphrodites for 180 additional neurons, grouped by anatomical region. Columns indicate the α, γ1, and γ2 reporters. Black circles indicate detection in all scored observations; gray circles indicate both positive and negative observations; open circles indicate no detection in any scored observation. Blank entries indicate that no usable detection calls were available. Asterisks identify negative-only assignments for which all recorded observations were obtained with the neuron on the side farther from the coverslip; these assignments may reflect reduced detectability.Sample sizes vary among neuron–reporter combinations; positive and negative observation counts are provided in Supplementary Table 1.

**Supplementary Figure 3.**
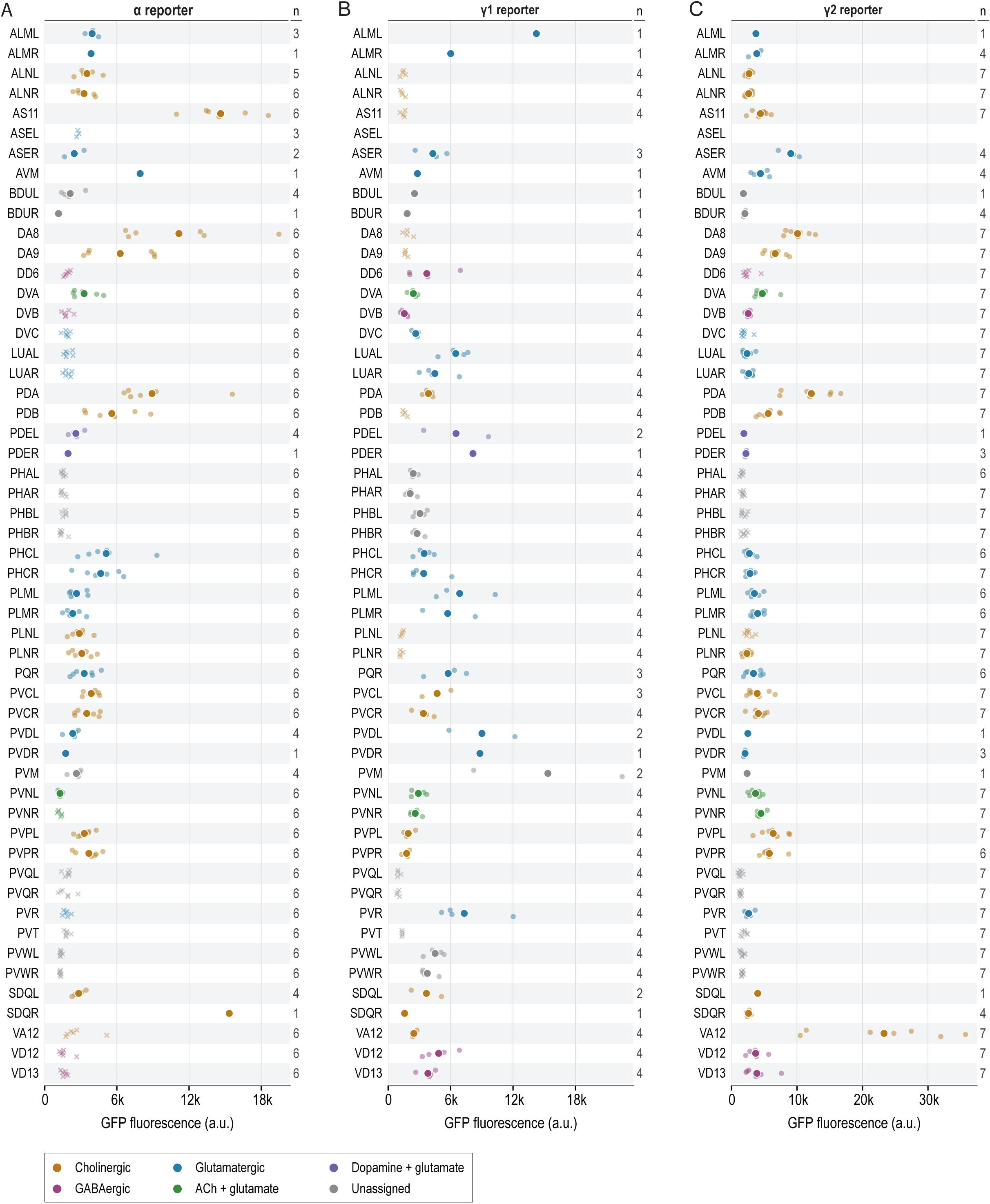
Fluorescence intensity of nrx-1 reporters in identified neurons. Estimated GFP fluorescence for the α **(A)**, γ1 **(B)**, and γ2 **(C)** reporters, with neurons aligned alphabetically across panels. Pale crosses show individual measurements for a neuron in which the reporter was scored as “not detected” in every L4 observation. For neurons with at least one positive call, small circles show all available intensity measurements, and larger circles with white borders indicate means. Blank rows indicate that no intensity measurement was available. Panels A, B, and C include 53, 52, and 52 neurons, comprising 266, 173, and 299 measurements, respectively. Numbers beside each row indicate the number of intensity measurements, which may differ from the number of observations used to score reporter detection in Figure 2 and Supplementary Figure 2. Fluorescence axis ranges differ across panels, and no additional background correction or normalization was applied during analysis. Absolute fluorescence values should not be directly compared across reporters. Measurements include observations qualitatively scored as reporter-negative. Colors indicate neurotransmitter identity, and gray indicates an unassigned classical neurotransmitter identity. Transmitter annotations follow Wang et al. (2024), Supplementary file 2; * indicates a potential glutamatergic assignment for PHA.

**Supplementary Figure 4.**
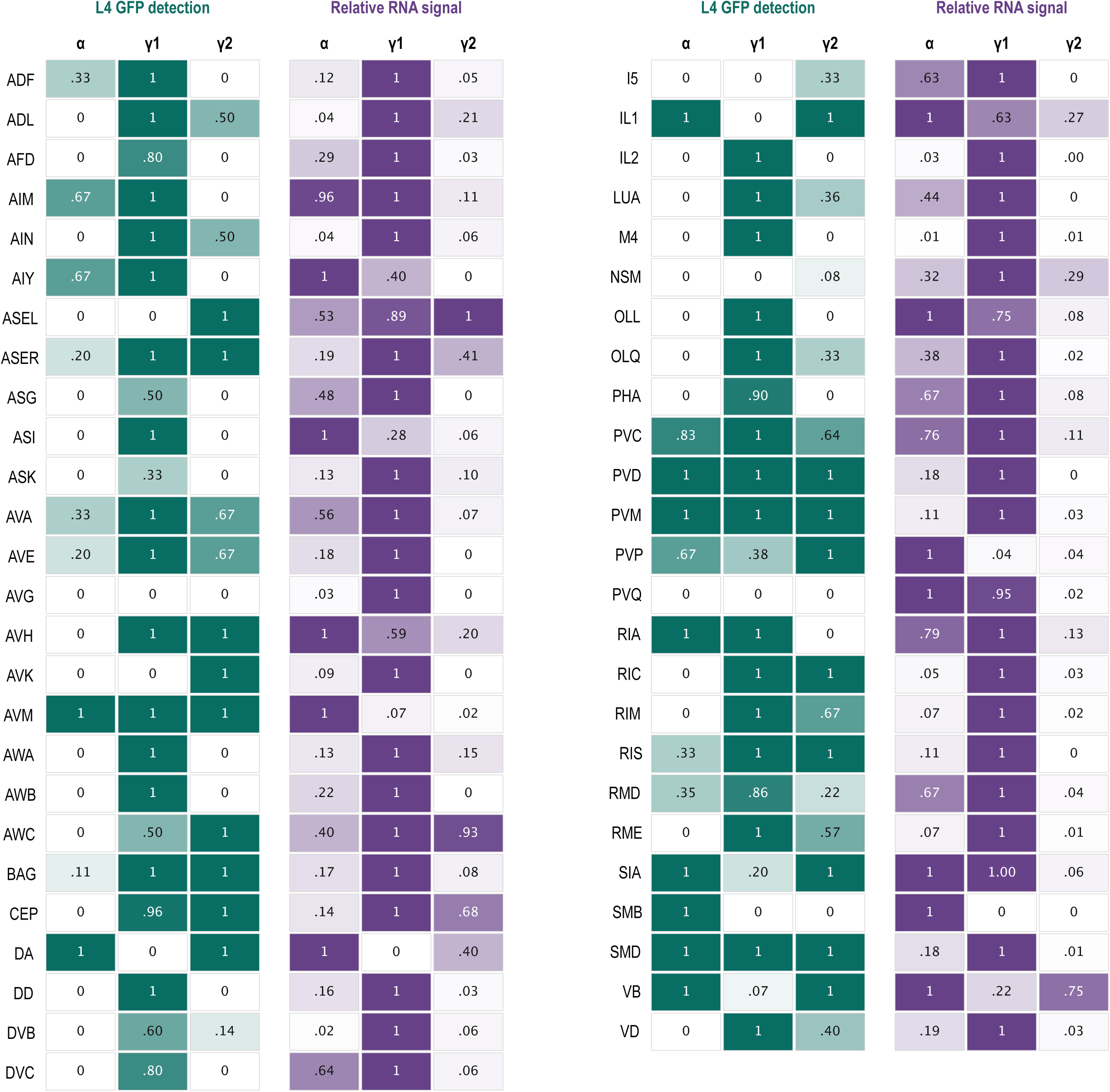
*nrx-1* reporter detection and relative RNA-seq signal across 51 neuron classes. Green cells show the fraction of scored L4 observations in which each reporter was detected in the NeuroPAL-identified neurons corresponding to each CeNGEN class. Purple cells show CeNGEN RNA signals for the three reporter groups, scaled so that the strongest signal within each class equals 1. The α signal reflects coverage at the α coding start. The γ1 signal reflects coverage at the *c/d* coding start after subtraction of α coverage to account for shared sequence. The γ2 signal reflects the local rise in coverage at the *nrx-1f* transcript start. Across neuron classes, Spearman correlations between GFP detection frequency and the plotted relative RNA signal were ρ = 0.43 for α, 0.33 for γ1, and 0.23 for γ2.

